# BioLogic, a parallel approach to cell-based logic gates

**DOI:** 10.1101/303032

**Authors:** Felipe A. Millacura, Brendan Largey, Christopher E. French

**Author notes:** Correspondence (F.M.); (C.F.).

## Abstract

In vivo logic gates have proven difficult to combine into larger devices. Our cell-based logic system, BioLogic, decomposes a large circuit into a collection of small subcircuits working in parallel, each subcircuit responding to a different combination of inputs. A final global output is then generated by a combination of the responses. Using BioLogic, for the first time a completely functional 3-bit full adder and full subtractor were generated using *Escherichia coli* cells; as well as a calculator-style display that shows a numeric result, from 0 to 7, when the proper 3 bit binary inputs are introduced into the system. BioLogic demonstrates the use of a parallel approach for the design of cell-based logic gates that facilitates the generation and analysis of complex processes, without the need for complex genetic engineering.

## Introduction

A major challenge in the field of synthetic biology is the construction of complex logic circuits that analyze variables as in electronics; where a single circuit accepts one or more binary inputs to generate one or more binary outputs. A cell-based logic network consists of engineered cells producing an output macromolecule only if the corresponding pattern of inputs is present. The mechanism of analysis is commonly based on the use of transcriptional regulators, transcription factors, polymerases, receptors or recombinases (Brenner et al., 2018). Some examples of genetic circuits mimicking computational behavior are toggle switches, oscillators, boolean logic gates, feedback controllers and multiplexers. Although there are genetic circuits that simulate computational behavior, the complex engineering of their biological chassis is affected by gene expression noise, mutation, cell death, undefined and changing extracellular environments and improper interactions with the cellular context (Andrianantoandro, et al., 2006). Furthermore, complex genetic engineering is necessary when multiple input variables are analyzed, limiting the processing capacity of the system.

Biological multiplexers analyze one or more signals over a common transmission line using interconnected transcription factors, recombinases, antisense RNA or CRISPR-like technology (Nielsen and Voight, 2014; Roquet et al., 2016; Brenner et al., 2018). However, complex genetic engineering is needed for wiring the basic computational units, becoming inefficient for moving beyond simple NOT or AND logic gates or for scaling to 3 bit logic circuits. The complexity of the genetic engineering required can be reduced by using distributed logic circuits, where the computation is distributed among several physically separated cellular consortia that each sense only one signal and respond by secreting a communication molecule (Regot et al., 2011). As a circuit responds to one signal, but not another, due to spatial distribution, a change in the state of the system can be triggered as response, making synthetic learning possible (Macia., et al 2017; Shipman et al., 2017). Even though the consortium approach makes Boolean circuit design simpler, it still shows a slow response and considerable complexity since each cell needs to recognize, synthesize and secrete a wiring molecule (Macia., et al 2016).

Here we propose an alternative logic architecture, which decomposes a large circuit into a collection of small subcircuits acting in parallel (hereafter BioLogic). Rather than having a single type of agent (such as a genetically engineered cell) doing the computation, BioLogic has separate types of agent that each react to a different combination of inputs. A final output is then generated by combination of the responses, making all kinds of binary operation possible. As an example, here we show the implementation of this concept using cells resistant to different combinations of antibiotics, with the response indicated by growth. This is used to demonstrate a completely functional 3 bit full adder and full subtractor, as well as a calculator-style display that shows digits from 0 to 7 based on three binary input bits.

## Methodology

### Reagents and stock solution preparations

Antibiotic stock solutions were prepared as follows: 100 mg/ml carbenicilin disodium salt (Sigma-Aldrich #C1389), 50 mg/ml kanamycin sulfate (PanReac Applichem #A1493), 20 mg/ml chloramphenicol (Acros Organics #22792), 10 mg/ml tetracycline hydrochloryde (Duchefa Biochemie #T0150), 10 mg/ml gentamicin sulfate (Melford #G0124), and 50 mg/ml spectinomycin.HCl (LKT Labs #S6018). Developing solution contained 0.1 %w/v bromothymol blue (Sigma-Aldrich #114421) and 400 mM Trizma base 400 mM pH7.5 (Sigma-Aldrich #T1503).

### Generation of subcircuit cells

*E. coli* JM109 was transformed with 200-300 pg of plasmid pSB4A5 (AmpR) or pSB4C5 (ChlR) (Registry of Standard Biological Parts) and selected on 100 µg/ml carbenicilin (Am) or 20 µg/ml chloramphenicol (Ch), respectively. Cells carrying the first bit plasmid were made chemically competent (Chung et al., 1989) and transformed with 200-300 pg of the 2nd bit plasmid, pSB1T3 (TetR) or pSB1K3 (KanR) (Registry of Standard Biological Parts). Selection was performed with the first antibiotic (Am or Ch) and the addition of 10 µg/ml Tetracycline (Tc) or 50 µg/ml kanamycin (Km), obtaining the two-bit combinations Km/Am (KA), Tc/Am (TA), Km/Ch (KC) and Tc/Ch (TC). This set of strains is sufficient to implement all two-bit binary operations.

The third bit layer was generated by transforming these four strains with pSEVA631 (GenR) (Silva-Rocha, et al., 2012) or pMO9075 (SpeR) (Keller, et al., 2011). Resulting strains were selected on the 2 bit antibiotic combinations plus 10 µg/ml gentamicin (Gm) or 50 µg/ml spectinomycin (Sm). This gave 8 strains, designated GTA (Gm/Tc/Am), GKA (Gm/Km/Am), STA (Sm/Tc/Am), SKA (Sm/Km/Am), GTC (Gm/Tc/Ch), STC (Sm/Tc/Ch), GKC (Gm/Km/Ch), SKC (Sm/Km/Ch) based on their resistance markers. This set of strains is sufficient to implement all three-bit binary operations. Plasmid specifications are listed in Table S1 and S3, with further information about these antibiotics in Table S2.

### Three-bit logic operations

Tests were performed in 96-well microplates by inoculating cells (1:100) in LB broth (100 uL) supplemented with 1%w/v D(+)-glucose (Fisher Chemical #G0500). Plates were incubated for 18 hours at 37°C without shaking and then developed by addition of the developing solution (0.1%w/v bromothymol blue in 400 mM Tris, pH7.5) in a ratio 1:20. Images were obtained using a Kodak ESPC315 Flatbed scanner. Design of the calculator-like display, full adder and subtractor are shown in Supplementary material (Figure S2 and S3).

### Results

In the distributed logic system of BioLogic, each input bit is has two forms, ZERO and ONE, each of which is essential to certain output agents and inhibitory to others. Thus each agent reacts only to a certain combination of input bits, allowing generation of any arbitrary pattern of outputs for any pattern of inputs. In the implementation shown here, each input bit comes in two forms, each being an antibiotic lethal to sensitive strains. In this case, bit A is represented by ampicillin for zero, chloramphenicol for one, bit B by kanamycin for zero, tetracycline for one, and bit C by spectinomycin for zero, streptomycin for one. Thus four strains are needed to implement any operation with two input bits, and eight strains for three input bits. In contrast to other cell-based logic schemes, only very minimal genetic engineering is required, essentially transformation with 3 different antibiotic resistance markers.

Cells show a global response concordant with the behavior expected for a 1 bit, 2 bit or 3 bit system (Figure 1). For instance, when the input 101 (chloramphenicol, tetracycline and spectinomycin) is added to the system growth is only observed in the corresponding STC cells, which carry the proper resistance markers. The response time of the system is around 12 hours (Figure S1) but plates were developed at 18 hours to avoid false negatives or positives.

**Figure 1:**
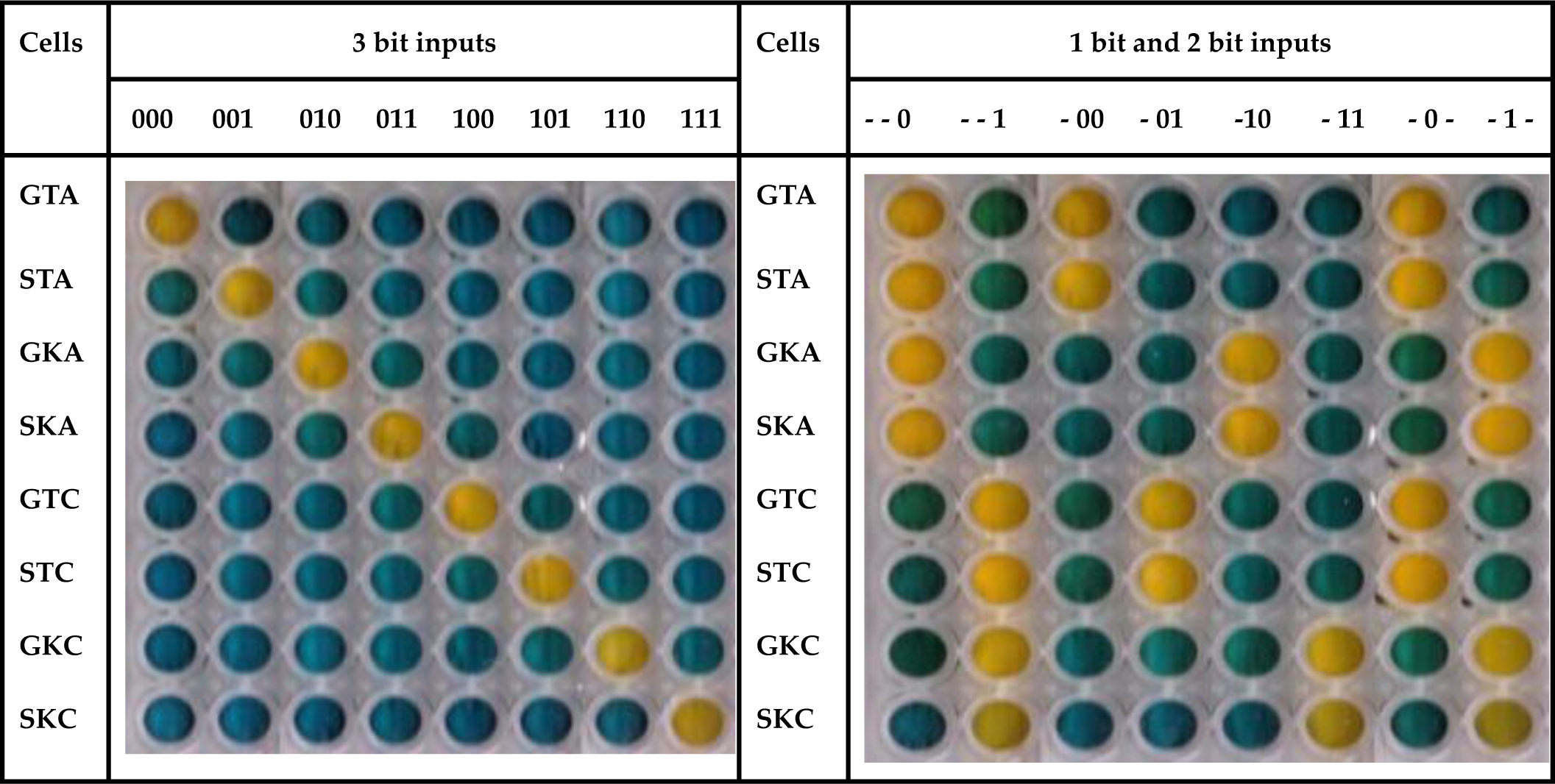
BioLogic responding to 1 bit, 2 bit and 3 bit inputs. BioLogic subcircuit cells were spatially distributed in different wells (vertically) and exposed to specified 1 bit, 2 bit or 3 bit inputs (top of each column). Cells were inoculated (1:100) in LB supplemented with 1% w/v glucose. After 18 hours of incubation at 37°C, plates were developed by addition of 0.05 volumes of the developing solution.

In order to further test the BioLogic system, a digital calculator-like display was designed (Figure S2). In this case, multiple subcircuit cells are mixed in one well and the global response displays a number from 0 to 7 when the proper binary input is applied. Therefore, when the input 110 represented by the antibiotics Gn/Km/Ch are added in the system, the number 6 is displayed (Figure 2).

**Figure 2.**
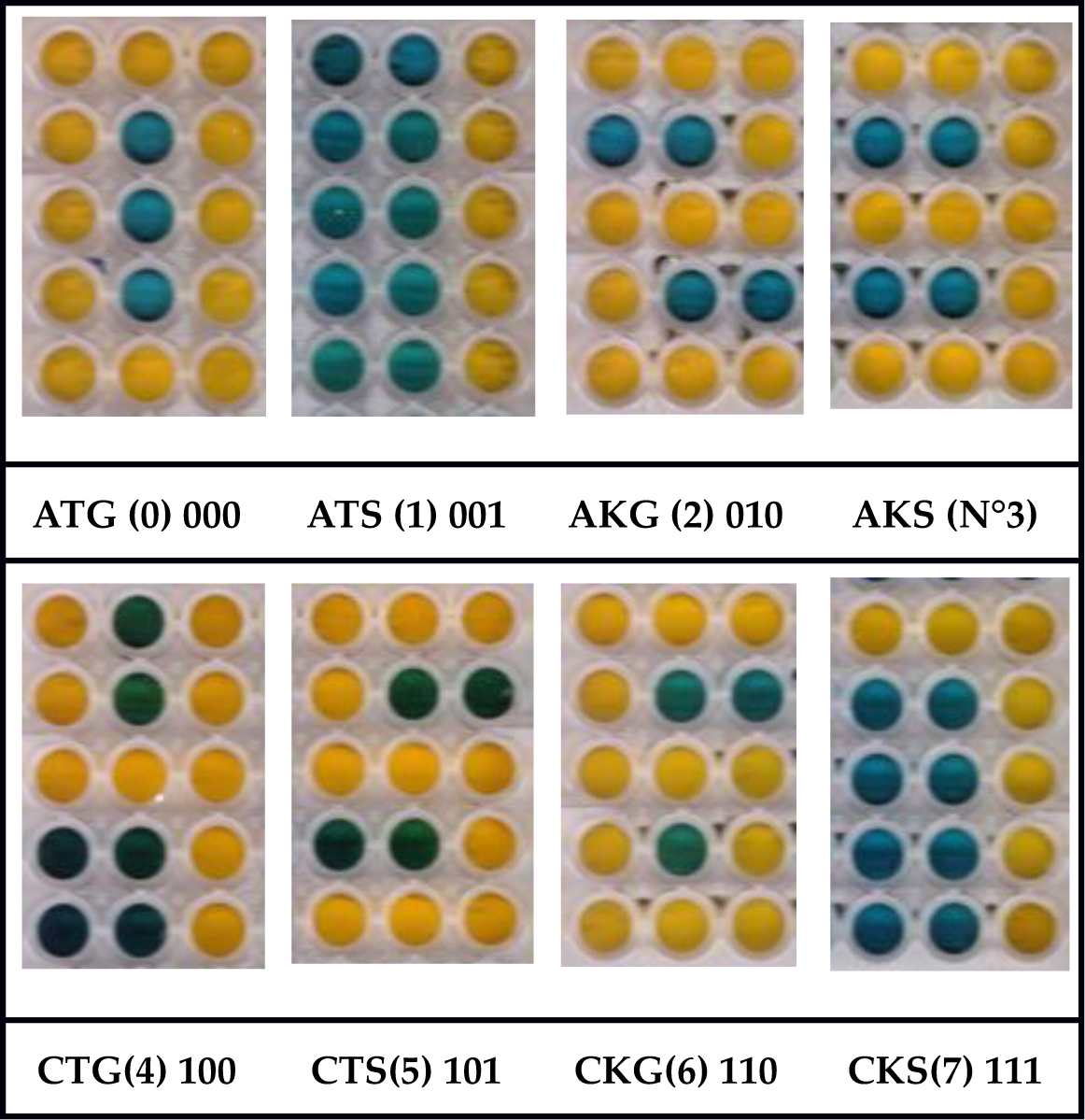
Digital calculator display using 3 bit BioLogic. The figure shows all numerals from zero to seven based on the 8 binary inputs provided. Cells were mixed as shown in Figure S2 and inoculated (1:100) in LB supplemented with 1%w/v glucose. After 18 hours of incubation, the plate was developed by addition of the 0.05 volumes of developing solution.

Finally, a full adder and a full subtractor were designed. Multiple subcircuit cells were mixed and distributed in two different wells (Figure S3). One well representing the solution (S) or difference (D) and a second one the carry (C_out_) or borrow (B_o_), for the adder and subtractor respectively (Figure 3).

**Figure 3.**
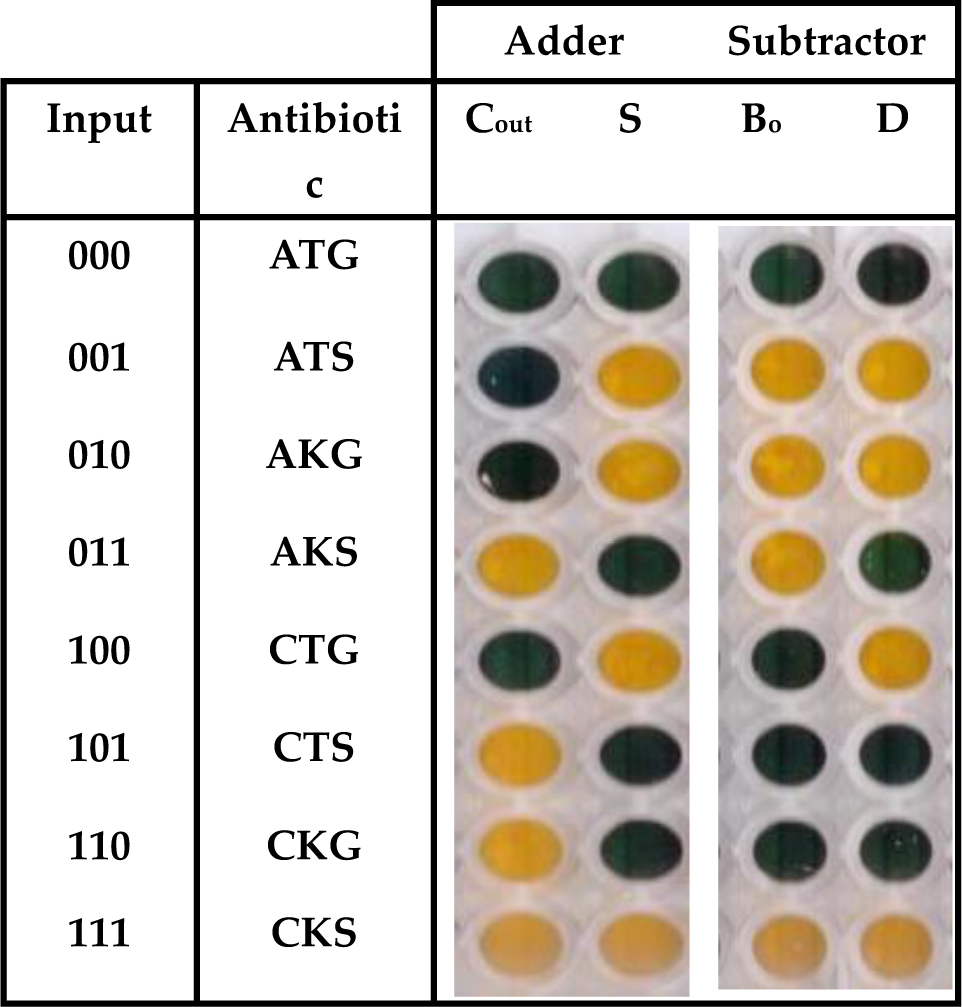
Full adder and subtractor using 3 bit BioLogic. The figure shows results of addition and subtraction using the BioLogic for 3 bit system. Cells were mixed as shown in Figure S3 and inoculated (1:100) in LB supplemented with 1%w/v glucose. After 18 hours of incubation, the plate was developed by addition of the 0.05 volumes of developing solution.

### Conclusions/Discussion

Subcircuits that solve complex calculations in parallel have been extensively used for computation in order to reduce the total computation time. Translating this approach to biological systems would allow us to analyze complex processes, currently difficult in synthetic biology, as multiple simple sub-circuits.

In our proof of concept, we present a biological information processing system, BioLogic, capable of exploiting the parallelism in mixed bacterial cultures. BioLogic decomposes the analysis of 2 and 3 bit complex inputs, into 4 and 8 sub circuits respectively (Figure 1). Each sub-circuit corresponds to a different *E. coli* strain carrying a different combination of antibiotic resistance markers (Table S1). As an example, in the 3 bit system the input 000 is represented by the antibiotics ampicillin, tetracycline and gentamicin (Figure 1). When this input is entered into the system, all cells that are not encoded for responding to 000 will die, but cells carrying the proper plasmid combination, pSB4A5, pSB1T3 and pSEVA621 will not (Figure 1 and 2), therefore, a live/dead response (output) is achieved in all sub circuits, the output of each well being one (growth) or zero (failure to grow) (Figure 2 and Figure S2).

BioLogic uses cellular consortia instead of a single type of cell. A similar approach has been developed by Macia et al. (2016 and 2017) using eukaryotic cells, and even showing the possibility of generating transient memory. However, that approach requires a sophisticated design as it relies on a secreted intermediate molecule (hormone-like) that must be kept at the right production level, and that should be previously activated by X (Repressor) and Y (ssrA-tagged protein) degradation. Furthermore, since the output of the circuit is distributed among different consortia, the concentration of the secreted molecule can differ according to the number of cells simultaneously producing it. This kind of multicellular approach and others based on single cells require sophisticated wiring design (Macia et al., 2016 and 2017; Siuti et al., 2013; Silva Rocha et al., 2008). By contrast, BioLogic requires very minimal genetic modification and little tuning to obtain reliable outputs (Figure 2 and 3).

The implementation of BioLogic presented here is simple, but its further development to useful applications presents a number of challenges; for example, expansion to 4 bits and beyond will require further well-behaved and non-cross-reacting antibiotic resistance markers, and will probably lead to even greater disparities in growth rate than those observed in the three-bit system (Figure S1). It will also be challenging to generate layered systems in which the output from one layer serves as input to another layer. However, the same concept, using a set of agents which each responds to a single combination of inputs, may also be implemented in other ways. One particularly attractive idea which we are pursuing is implementation in a cell-free transcriptional system, in which inputs may be present as small molecules interacting with transcription factors, or as DNA or RNA oligonucleotides; this will also allow transcriptional outputs from one layer to be used as inputs to a second layer. By this means we hope to generate complex binary logic systems which may be used in a variety of synthetic biology applications.

## Supporting information

## Supplementary Materials

The following are available in the Supplementary section. Figure S1. BioLogic cell growth curves. Figure S2: BioLogic calculator-like display design. Figure S3: BioLogic 3 bit Full adder/subtractor design. Table S1: Plasmids used for generating BioLogic subcircuit strains. Table S2: Antibiotics and resistance cassettes used on BioLogic. Table S3: Plasmid incompatibility groups.

## Author Contributions

F.A.M., and C.F.. conceived and designed the experiments; F.A.M., and B.L. performed the experiments; F.A.M. and C.F. analysed and validated the data; F.A.M and C.F. contributed reagents/materials/analysis tools; F.A.M. and C.F.. wrote – reviewed & edited the manuscript.

## Acknowledgments

The authors acknowledge the valuable assistance of Dr. Louise Horsfall for supplying the pMO9075 plasmid and Dr. Aitor De las Heras for supplying the pSEVA623 plasmid. The authors also acknowledge the following funding source: CONICYT/BC-PhD 72170403 (FM)

---

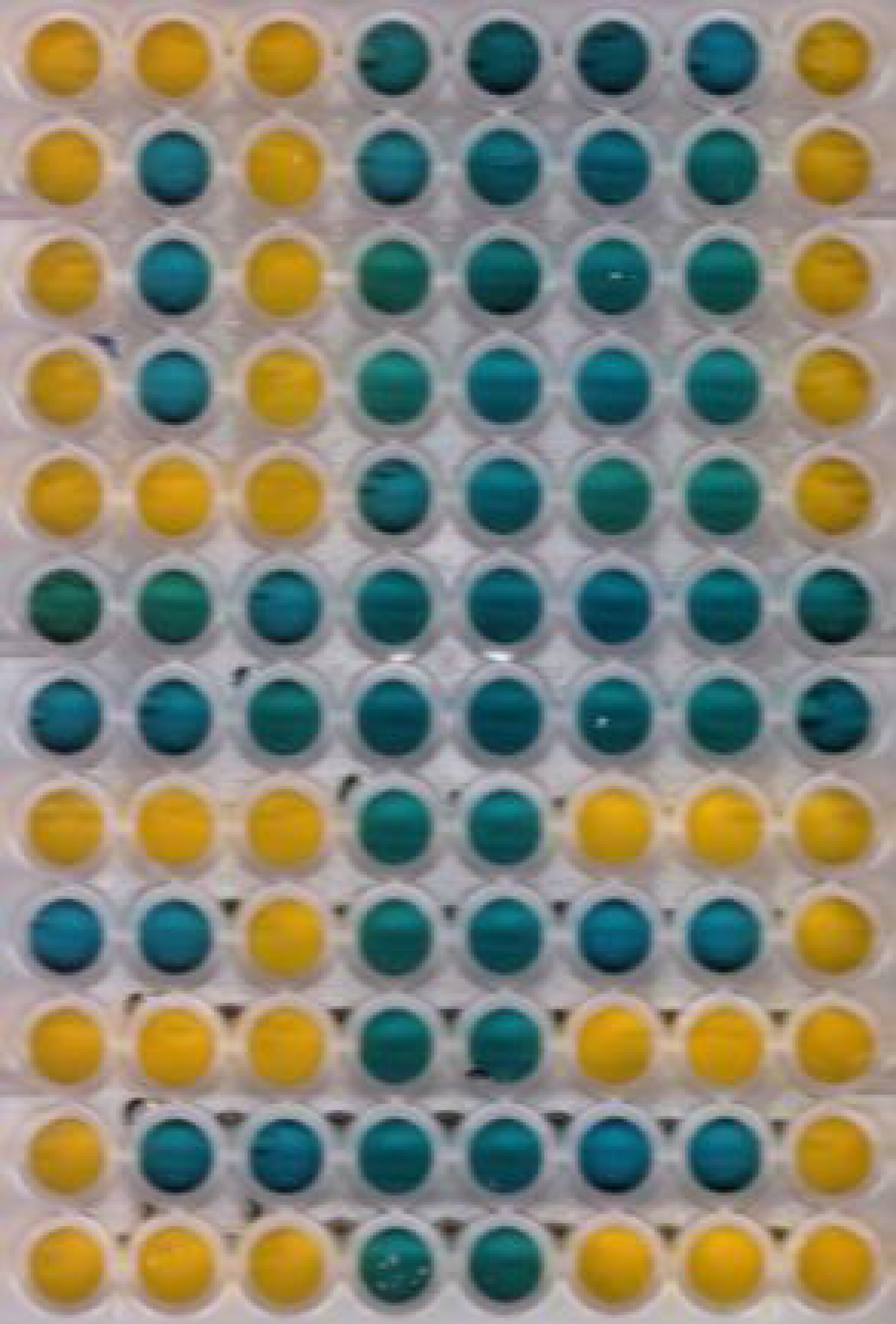

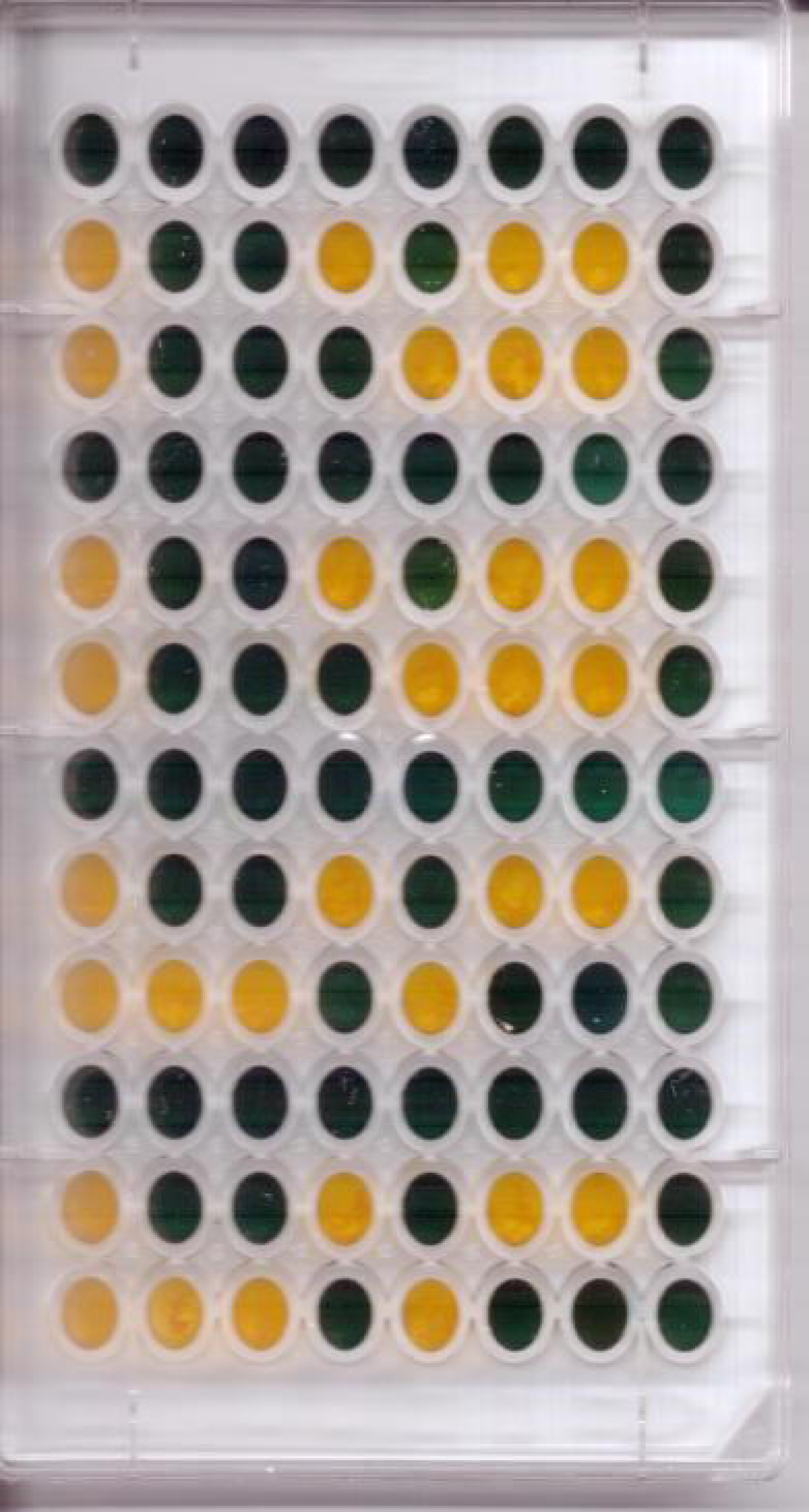

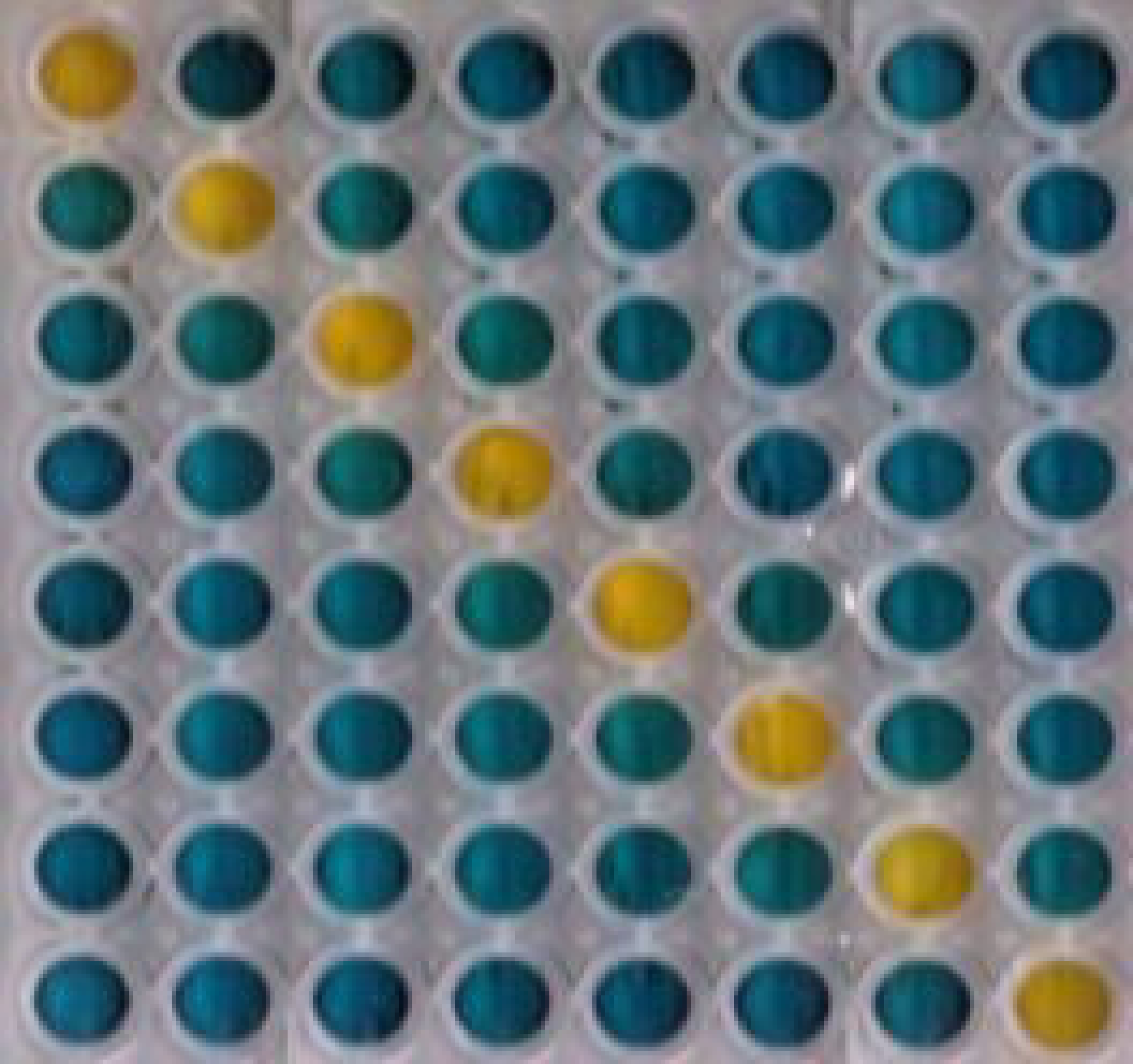

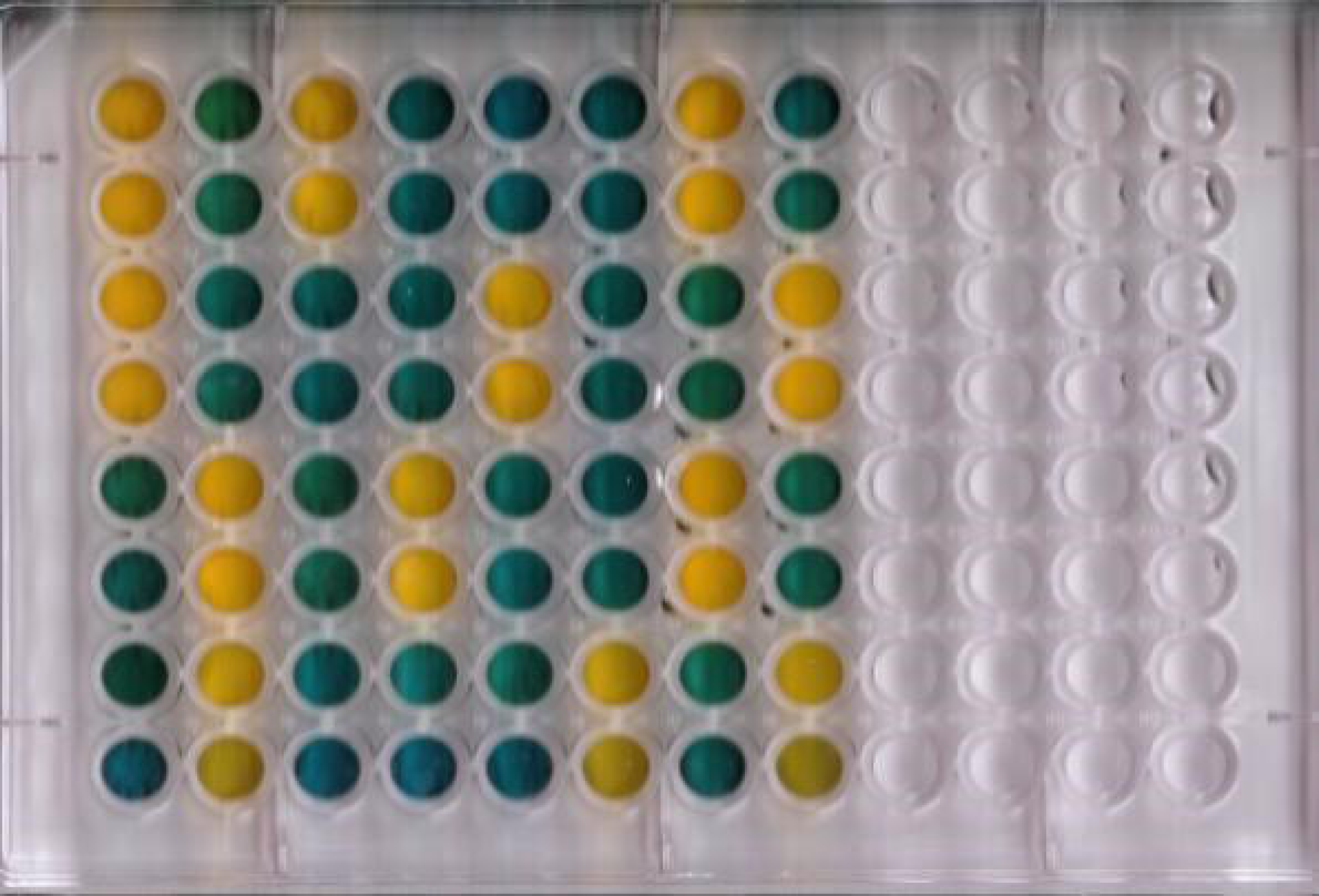

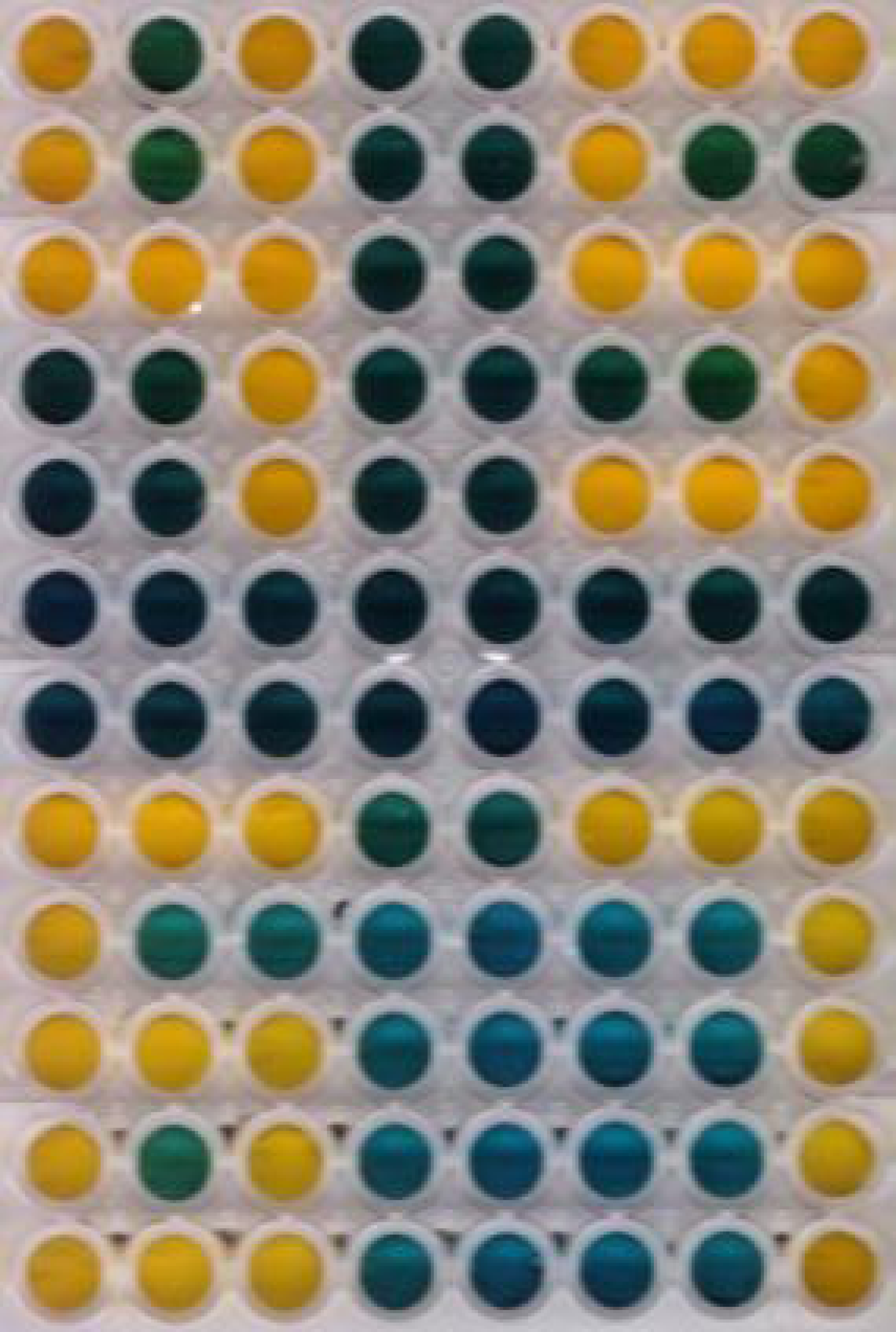

## References

1. Andrianantoandro, E., Basu, S., Karig, D. K., & Weiss, R. (2006). Synthetic biology: new engineering rules for an emerging discipline. Molecular systems biology, 2(1).

2. Brenner, M. J., Cho, J. H., Wong, N. M., & Wong, W. W. (2018). Synthetic Biology: Immunotherapy by Design. Annual review of biomedical engineering, (0).

3. Begbie S. et al., (2005) The Effects of Sub-Inhibitory Levels of Chloramphenicol on pBR322 Plasmid Copy Number in *Escherichia coli* DH5a. Journal of Experimental Microbiology and Immunology (JEMI). 7:82–8.

4. Maniatis, T., Fritsch, E. F. and Sambrook, J. Molecular cloning. New York: Cold Spring Harbor Laboratory; 1982. pp 545.

5. Nielsen, A. A., & Voigt, C. A. (2014). Multi-input CRISPR/Cas genetic circuits that interface host regulatory networks. Molecular systems biology, 10(11), 763.

6. Chung, C. T., Niemela, S. L., & Miller, R. H. (1989). One-step preparation of competent Escherichia coli: transformation and storage of bacterial cells in the same solution. Proceedings of the National Academy of Sciences, 86(7), 2172–2175.

7. Silva-Rocha, R., Martínez-García, E., Calles, B., Chavarría, M., Arce-Rodríguez, A., de Las Heras, A.,… & Platero, R. (2012). The Standard European Vector Architecture (SEVA): a coherent platform for the analysis and deployment of complex prokaryotic phenotypes. Nucleic acids research, 41(D1), D666–D675.

8. Keller, K. L., Wall, J. D., & Chhabra, S. (2011). Methods for engineering sulfate reducing bacteria of the genus Desulfovibrio. In Methods in enzymology (Vol. 497, pp. 503–517). Academic Press.

9. Macia J, Vidiella B, Solé RV. 2017 Synthetic associative learning in engineered multicellular consortia. J. R. Soc. Interface 14: 20170158.

10. Macia, J., Manzoni, R., Conde, N., Urrios, A., de Nadal, E., Solé, R., & Posas, F. (2016). Implementation of complex biological logic circuits using spatially distributed multicellular consortia. PLoS computational biology, 12(2), e1004685.

11. Roquet, N., Soleimany, A. P., Ferris, A. C., Aaronson, S., & Lu, T. K. (2016). Synthetic recombinase-based state machines in living cells. Science, 353(6297), aad8559.

12. Regot, S., Macia, J., Conde, N., Furukawa, K., Kjellén, J., Peeters, T., … & Solé, R. (2011). Distributed biological computation with multicellular engineered networks. Nature, 469(7329), 207.

13. Shipman, S. L., Nivala, J., Macklis, J. D., & Church, G. M. (2017). CRISPR–Cas encoding of a digital movie into the genomes of a population of living bacteria. Nature, 547(7663), 3

