## Supplementary material for "BioLogic, a parallel approach to cell-based logic gates"

### Supplementary data

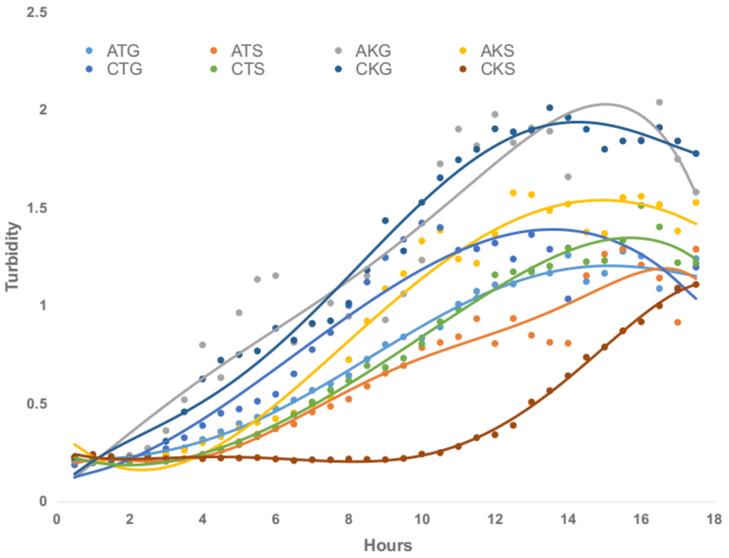


**Figure S1. BioLogic cell growth curves.** Growth curves of the 3-bit subcircuit cells in LB + 0.1% glucose with their respective antibiotic combinations. **A:** Carbenicilin (100 µg/mL), **C:** Chloramphenicol (20 µg/mL) **T:** Tetracycline (10 µg/mL), **K:** Kanamycin (50 µg/mL), **G:** Gentamicin (10 µg/mL). **S:** Spectinomycin (50 µg/mL). Overnight culture (0.01 volume) was used as inoculum.

**Figure S2: BioLogic calculator-like display design**

#### Cell distribution

| | | |
|---|---|---|
| A | B | C |
| D | --- | E |
| F | G | H |
| I | --- | J |
| K | L | M |

| No. to display | A | B | C | D | E | F | G | H | I | J | K | L | M |
|---|---|---|---|---|---|---|---|---|---|---|---|---|---|
| **000 (0)** | ATG | ATG | ATG | ATG | ATG | ATG | ---- | ATG | ATG | ATG | ATG | ATG | ATG |
| **001 (1)** | ---- | ---- | ATS | ---- | ATS | ---- | ---- | ATS | ---- | ATS | ---- | ---- | ATS |
| **010 (2)** | AKG | AKG | AKG | ---- | AKG | AKG | AKG | AKG | AKG | ---- | AKG | AKG | AKG |
| **011 (3)** | AKS | AKS | AKS | ---- | AKS | AKS | AKS | AKS | ---- | AKS | AKS | AKS | AKS |
| **100 (4)** | CTG | ---- | CTG | CTG | CTG | CTG | CTG | CTG | ---- | CTG | ---- | ---- | CTG |
| **101 (5)** | CTS | CTS | CTS | CTS | ---- | CTS | CTS | CTS | ---- | CTS | CTS | CTS | CTS |
| **110 (6)** | CKG | CKG | CKG | CKG | ---- | CKG | CKG | CKG | CKG | CKG | CKG | CKG | CKG |
| **111 (7)** | CKS | CKS | CKS | ---- | CKS | ---- | ---- | CKS | ---- | CKS | ---- | ---- | CKS |

Cells added in specified wells carry resistance markers for:

**ATG:** Carbenicilin, Tetracycline, Gentamicin.

**ATS:** Carbenicilin, Tetracycline, Spectinomycin.

**AKG:** Carbenicillin, Kanamycin, Gentamicin.

**AKS:** Carbenicillin, Kanamycin, Spectinomycin.

**CTG:** Chloramphenicol, Tetracycline, Gentamicin.

**CTS:** Chloramphenicol, Tetracycline, Spectinomycin.

**CKG:** Chloramphenicol, Kanamycin, Gentamicin.

**CKS:** Chloramphenicol, Kanamycin, Spectinomycin.

**Figure S3: BioLogic 3 bit Full adder/subtractor design**

| | | Adder | | Subtractor | |
|---|---|---|---|---|---|
| **Input** | **Antibiotic** | $C_{out}$ | **S** | $B_o$ | **D** |
| **000** | **ATG** | ---- | ---- | ---- | ---- |
| **001** | **ATS** | ---- | +++ | +++ | +++ |
| **010** | **AKG** | ---- | +++ | +++ | +++ |
| **011** | **AKS** | +++ | ---- | +++ | ---- |
| **100** | **CTG** | ---- | +++ | ---- | +++ |
| **101** | **CTS** | +++ | ---- | ---- | ---- |
| **110** | **CKG** | +++ | ---- | ---- | ---- |
| **111** | **CKS** | +++ | +++ | +++ | +++ |

Cells added in specified wells carry resistance markers for:

**ATG:** Carbenicilin, Tetracycline, Gentamicin.

**ATS:** Carbenicilin, Tetracycline, Spectinomycin.

**AKG:** Carbenicillin, Kanamycin, Gentamicin.

**AKS:** Carbenicillin, Kanamycin, Spectinomycin.

**CTG:** Chloramphenicol, Tetracycline, Gentamicin.

**CTS:** Chloramphenicol, Tetracycline, Spectinomycin.

**CKG:** Chloramphenicol, Kanamycin, Gentamicin.

**CKS:** Chloramphenicol, Kanamycin, Spectinomycin.

**Additional information**

**Table S1: Plasmids used for generating BioLogic subcircuit strains.**

| Plasmid | Antibiotic marker | ORI | Copy number | Reference |
|---|---|---|---|---|
| pSB4A5 | AmpR | pSC101 | 5 | http://parts.igem.org/Part:pSB4A5 |
| pSB4C5 | CmlR | pSC101 | 5 | http://parts.igem.org/Part:pSB4C5 |
| pSB1T3 | TetR | pMB1(der) | 100-300 | http://parts.igem.org/Part:pSB1T3 |
| pSB1K3 | KanR | pMB1 (der) | 100-300 | http://parts.igem.org/Part:pSB1K3 |
| pSEVA631 | GenR | pBBR1 | medium | Silva-Rocha et al., 2013 |
| pSEVA621 | GenR | trfA | low | Silva-Rocha et al., 2013 |
| pMO9075 | SmR | pBG1 | low | Keller, et al., 2011 |

**Table S2: Antibiotics and resistance cassettes used on BioLogic.**

| Antiobiotic | Class | Mode of action | Resistance |
|---|---|---|---|
| Carbenicilin | β-lactam | Bactericidal; Inhibits cell wall synthesis | β-lactamase (bla) gene |
| Kanamycin | Aminoglycoside | Bactericidal; Binds 30S ribosomal subunit; causes mistranslation | Neomycin phosphotransferase II |
| Cloramphenicol | Chloramphenicol | Bacteriostatic; Binds 50S ribosomal subunit; inhibits peptidyl translocation | Chloramphenicol acetyl transferase |
| Tetracycline | Tetracycline | Bacteriostatic; Binds 16S ribosomal subunit; inhibits protein synthesis (elongation step) | Tetracycline efflux protein |
| Gentamicin | Aminoglycoside | Irreversibly binding the 30S subunit of the bacterial ribosome | Gentamicin-3-N-acetyltransferase |
| Spectinomycin | Aminocyclitol | It binds to the 30S and interrupts protein synthesis affecting 16S rRNA | Spectinomycin adenyltransferase |

**Table S3: Plasmid incompatibility groups.**

| Incompatibility group | Regulation | Comment |
|---|---|---|
| pBR322/ColE1/pMB1 | Inhibitor-target RNA I | Control processing of pre-RNAII into primer |
| IncFII, pT181 | RNA | Affecting synthesis of RepA protein |
| R6K*, P1, F, pSC101, Rts1, P15A*, RK2 | Iteron binding | Sequestering of RepA protein |
